# Evolution of gene order in prokaryotes is driven primarily by gene gain and loss

**DOI:** 10.1101/2025.04.03.647019

**Authors:** Shelly Brezner, Sofya K. Garushyants, Yuri I. Wolf, Eugene V. Koonin, Sagi Snir

## Abstract

Evolution of bacterial and archaeal genomes is highly dynamic including extensive gene gain via horizontal gene transfer and gene loss as well as different types of genome rearrangements, such as inversions and translocations, so that gene order is not highly conserved even among closely related organisms. We sought to quantify the contributions of different genome dynamics processes to the evolution of the gene order relying on the recently developed “jump” model of gene translocation. The jump model has been completely solved analytically and provides the exact distribution of syntenic gene block lengths (SBL) in compared genomes based on gene translocations alone. Comparing the SBL distribution predicted by the jump model with the distributions empirically observed for multiple groups of closely related bacterial and archaeal genomes, we obtained robust estimates of the genome rearrangement to gene flux (gain and loss) ratio. In most groups of bacteria and archaea, this ratio was found to be on the order of 0.1 indicating that the loss of synteny in the evolution of bacteria and archaea is driven primarily by gene gain and loss rather than by gene translocation.

**Significance:** Evolution of bacterial and archaeal genomes is a highly dynamic process that includes extensive gene gain via horizontal gene transfer and gene loss as well as different types of genome rearrangements, so that gene order is not highly conserved even among closely related organisms. We developed a theoretical framework to quantify the contributions of different genome dynamics processes to the evolution of the gene order and found that in most groups of bacteria and archaea, the genome rearrangement to gene flux (combined gain and loss) is on the order of 0.1. Thus, the loss of genomic synteny in the evolution of bacteria and archaea appears to be driven primarily by gene gain and loss rather than by gene translocation.

## Introduction

Evolution of bacterial and archaeal (collectively, prokaryote) genomes is a highly dynamic process that, in addition to the accumulation of point mutations and small, within gene, indels, involves extensive gene loss and gene gain, primarily, via horizontal gene transfer (HGT) (1-7). Multiple, independent studies on prokaryote genome evolution consistently show that gene gain and loss (to which we collectively refer as gene flux) occur at rates about an order of magnitude higher than that of gene duplication indicating that gene flux is central to prokaryote evolution (8-10).

One of the well-known, prominent manifestations of the dynamic evolution of prokaryote genomes is the lack of conservation of the gene order (genomic synteny) that in many cases is observed even between closely related bacteria and archaea (3, 11-14). Functionally linked prokaryote genes are often organized into operons, arrays of (typically) 2-5 genes that are co-transcribed into a single RNA molecule ensuring coordination of expression of these genes (13, 15-17). Many operons are shared by genomes of diverse bacteria and archaea, in part, because HGT of complete operons appears to be favored by selection (16, 18-21). However, on the scale above operons, gene synteny appears to deteriorate quickly during prokaryote evolution albeit apparently at broadly varying rates among different groups of bacteria and archaea (3, 13, 14).

In addition to gene loss and HGT, genome dynamics involves translocations and inversions of genome segments (22-24). Evidently, genomic synteny can be disrupted by different types of events including both gene flux and translocations and inversions. Our previous analysis of archaeal genome evolution showed that, although gene flux alone is insufficient to explain the observed decay of genome synteny, the propensity of a gene to be involved in genome rearrangement is proportional to its propensity for involvement in gene flux, suggesting a common basis for genome dynamics (25).

Recently, a theoretical model of gene order evolution as a Poisson random process (hereafter, “jump model”) has been developed and completely solved analytically (26). The jump model provides the distribution of the expected lengths of syntenic (gene) blocks lengths (SBL) in two compared genomes given the number of single gene translocations (jumps) since the divergence of the respective organisms from their common ancestor. Here, we took advantage of the jump model and comparisons of gene content and gene order in multiple groups of closely related bacterial and archaeal genomes to estimate the genome rearrangement to gene flux ratio for each group. We found that in most cases, the rate of gene flux is about an order of magnitude higher than the rate of genome rearrangement and thus gene flux makes the principal contribution to the decay of genome synteny in prokaryote evolution, at least, at short evolutionary distances.

## Results

### Comparison of gene content and gene order in bacterial and archaeal genomes

We used 172 sets of closely related bacterial and archaeal genomes (Alignable Tight Genome Clusters, ATGC, (27), AdditionalData_1) to investigate short-range evolution of gene content and gene order. High-resolution, ATGC specific COGs (clusters or orthologous genes, (27, 28)) were used to label genes in the genomes within each ATGC. We sought to analyze evolution of gene content and gene order in each ATGC in a mutually independent manner. To this end, gene content differences in a pair of genomes was measured as gene content distance (GCD), the log-corrected fraction of shared genes; see Methods for details), ignoring gene arrangement in the genome and effects of paralogy. Gene order conservation was analyzed by identifying all syntenic gene blocks in a pair of genomes (including “blocks” of length 1 and ignoring the gene coding directions) across the shared gene complement (thus, ignoring the differences between the gene sets resulting from gene flux). The distributions of SBL provided a comprehensive quantitative picture of synteny, allowing for nuanced comparisons between the observed data and the model predictions, whereas the mean SBL can serve as a single number characterizing the gene order similarity.

As expected, synteny and gene content are highly correlated in most of the ATGCs across all explored parts of the prokaryotic tree of life (Spearman rank correlation between GCD and mean SBL of -0.80 across all genome pairs within the 172 ATGCs). Nevertheless, there are substantial differences between the relative rates of gene exchange and genome shuffling among the ATGCs. For example, at the same level of GCD (∼0.01 gains/losses per gene), genomes of ATGC030 (*Mycobacterium abscessus*/*immunogenum*) retain enough synteny within the common gene subset to have the average SBL of ∼500, whereas genomes of ATGC183 (*Bordetella pertussis*/*bronchiseptica*/*parapertussis*) are shuffled enough to retain syntenic blocks of only 20-30 genes on average (Figure 1_distplot).

**Figure 1.**
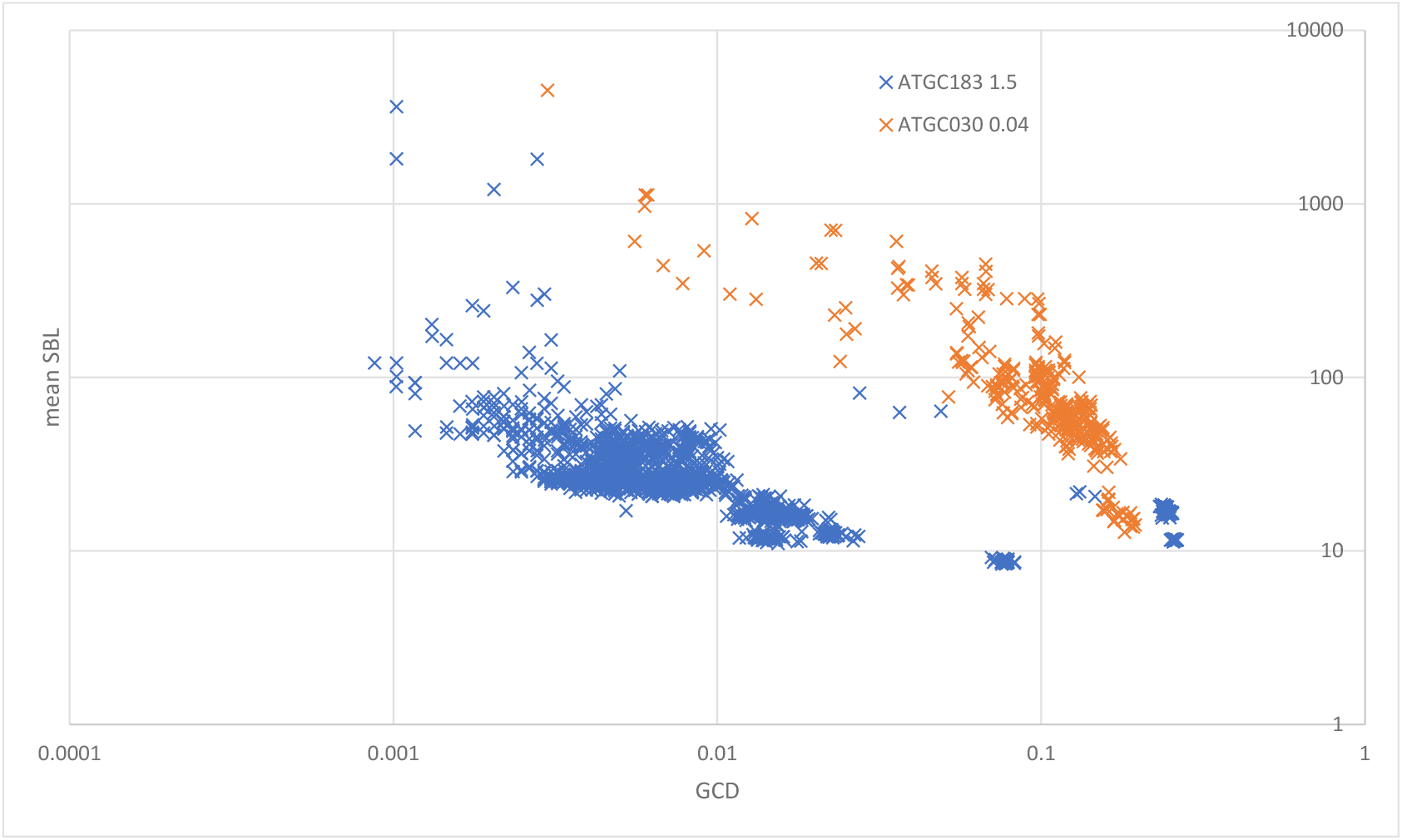
Mean Synteny Block Length vs Gene Content Distance. X-axis, gene contend distance (GCD). Y axis, mean synteny block length (SBL). Each data point represent the pair of genomes within an ATGC (blue, ATGC183, *Bordetella*; orange, ATGC030, *Mycobacterium*).

### Rearrangement to gene flux ratio

To quantify the relationships between the gene flux and gene shuffling we took advantage of the “jump model” (26) that was completely solved analytically. Tractable change of gene order in genomes that evolve by random single-gene translocations (“jumps”) that are equiprobable with respect to both the choice of the translocating gene and the destination, allows one to derive the precise distribution of SBL after the specified number of translocations.

The SBL distribution for a given pair of genomes depends on the number of genome rearrangements on the evolutionary trajectory between them. This number, in turn, depends on the evolutionary distance and the rate of rearrangement. Our analysis is based on two assumptions concerning this process: i) gene flux (loss and gain) rate is approximately constant within an ATGC and, therefore, gene content distance approximates a local, clade-specific clock and ii) genome rearrangement rate is approximately constant within an ATGC. The corollary of these two assumptions is that there exists an ATGC-specific rearrangement-to-flux ratio, that is, the characteristic number of rearrangements per gene gain or loss event.

For each pair of genomes *A* and *B* within an ATGC, the GCD *d*_*A,B*_ and the SBL distribution 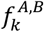 are computed from genomic data. Assuming an ATGC-specific ratio *q* between the number of jumps per gene *λ*_*A,B*_ and the number of gene gains and losses *d*_*A,B*_ (formally, *λ*_*A,B*_ = *qd*_*A,B*_), the theoretical SBL distribution 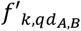 can be calculated for any given values of *q*. The difference between the observed distribution 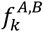 and the predicted distribution 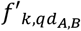 can be quantified using the Wasserstein distance metric 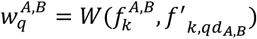, where *W*(*P, Q*) is 1-Wasserstein distance for the two distributions, *P* and *Q* (29). (Figure 2_SBLcomparison). Summing over all genome pairs within an ATGC obtains the global measure of the goodness of fit between the observed and expected SBL distributions for the given value of q, 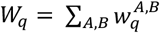. Minimization of *W*_*q*_ over *q* yields *q*^∗^, the ATGC-specific optimal value of the rearrangement-to-flux ratio.

**Figure 2.**
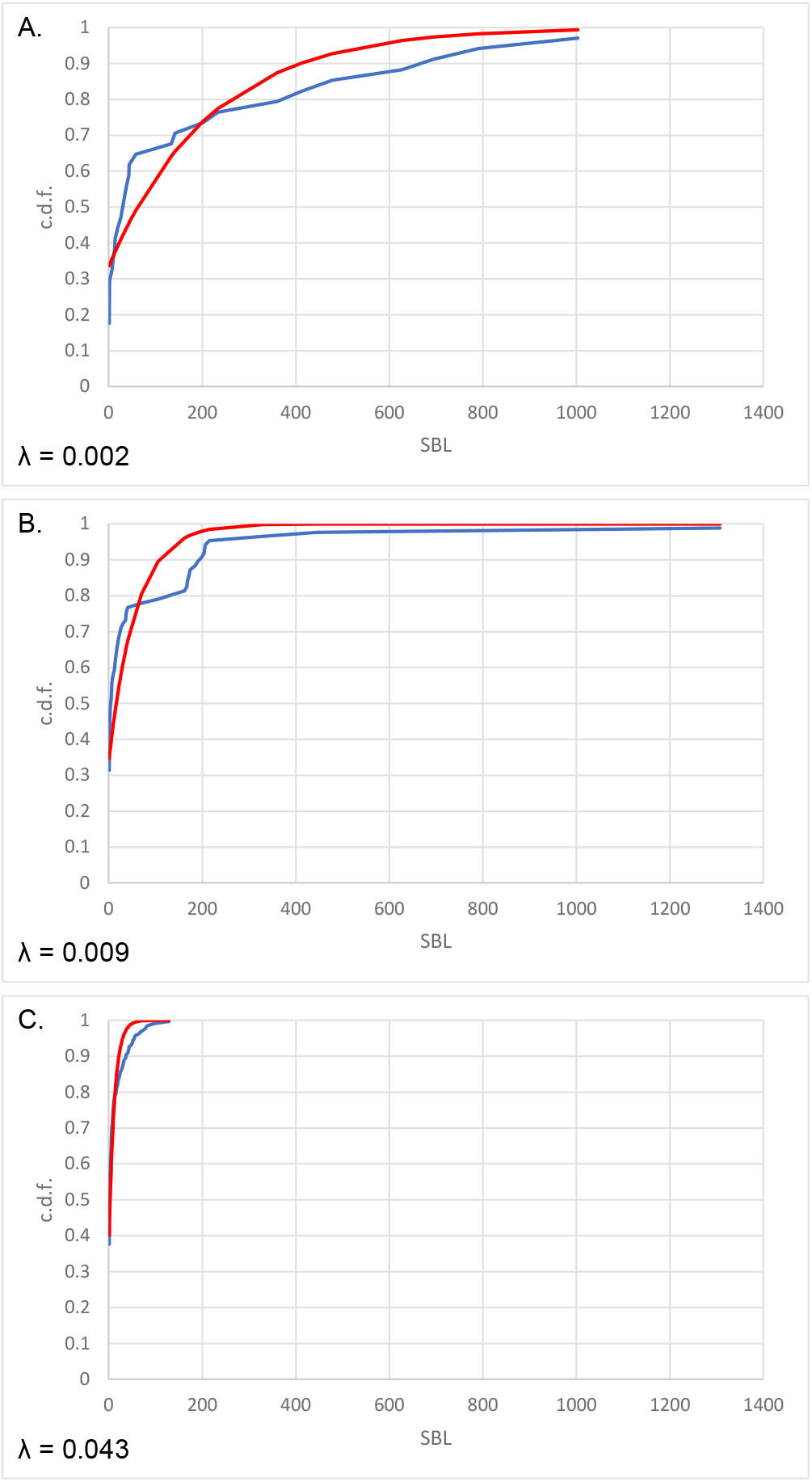
Fitting syntenic gene block distributions for ATGCs to the theoretical distribution given by the jump model. Blue: observed cumulative density function. Red: cumulative density function obtained from the jump model with the specified value of *λ* (ATGC002). A. *Klebsiella pneumoniae* UHKPC07 (GCF_000417265.2) vs *K. pneumoniae* UHKPC33 GCF_000417085.2 B. *K. pneumoniae* UHKPC07 (GCF_000417265.2) vs *K. pneumoniae* GCF_000775395.1 C. *K. pneumoniae* UHKPC07 (GCF_000417265.2) vs *K. pneumoniae* GCF_000784945.1

Robustness of the *q*^∗^ estimate was assessed by generating bootstrap samples of genome pairs and optimizing the value of *q* for each sample. We considered *q*^∗^ to be well-defined for a given ATGC if the values of the 5th and the 95th percentiles of the distribution of bootstrapped estimates differed by no more than a factor of 2.

For 139 of the 172 ATGCs, a robust estimate of the rearrangement to gene flux ratio was obtained, operationally defined as the 6th and 95th ranked values among the 100 bootstrap samples (middle 90% of the range) differing by no more than a factor of 2 (AdditionalData_2). Among these 139 ATGc, the values of *q*^∗^ covered the range from 0.04 to 2.5, with the median of 0.13 and the value of 1 exceeded for only 5 ATGCs (Figure 3_qdistr).

**Figure 3.**
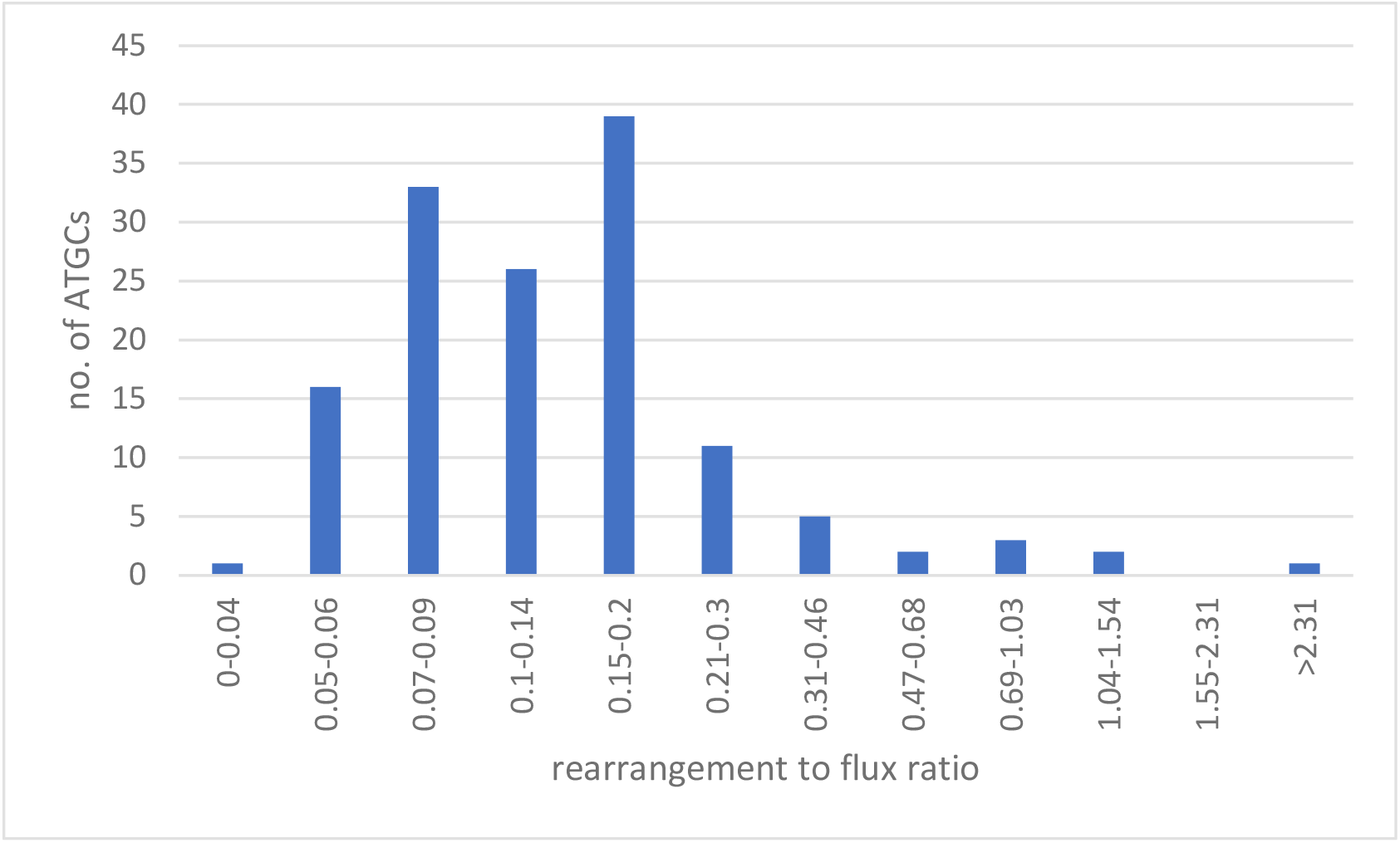
Distribution of the rearrangement to gene flux ratios across the ATGCs. X-axis, rearrangement to flux ratio (*q*^∗^), logarithmically binned. Y-axis, number of ATGCs.

If the gains and losses are balanced, preserving the genome size, the effect of the gene flux on synteny can be assessed by considering pairs of gene gains and losses. When the number of events is small relative to the genome size, the effect of a single random gene translocation is the same as the effect of a single random loss-gain pair (both involve one gene leaving its original location and one gene appearing in the new location, the only difference being that, in the loss-gain case, the gene at the new location is likely to have never been present in this genome before). Thus, in most ATGCs, there are 5-13 gain-loss pairs per translocation and, therefore, 85-95% of the synteny disruptions can be accounted for by gene flux.

### Patterns of relative rates of gene rearrangement and gene flux

Neither the well-defined values of *q*^∗^nor the pattern of which ATGCs did and which did not yield well-defined values, showed any obvious association with gross characteristics of ATGCs such as number of genomes, phylogenetic depth, genome size and others; AdditionalData_1, AdditionalData_2). The phylogenetic relatedness between ATGCs was found to be only weakly correlated with the rearrangement-to-flux ratio (*R*^2^ 0.008, p-value 0.27, SI Figure S1), with large differences in *q*^∗^ values observed for some closely related ATGCs. For example, *Pseudomonas, Bordetella* and *Burkholderia* ATGCs span the range of *q*^∗^ values of more than an order of magnitude within a genus (AdditionalData_2, SI Figure S2).

In terms of presence or absence of specific genes, no single COG or COG functional category could account for the differences in the rearrangement to flux ratios. In particular, no correlation was detected between the *q*^∗^ values and the presence or absence of genes implicated in intra-genomic recombination, such as those involved in non-homologous end joining (30) (AdditionalData_3). Likewise, phylogenetic correlation failed to identify a set of COGs that would explain the differences in the rearrangement to gene flux ratios between ATGCs (AdditionalData_4_PGLS). Both the incremental (adding COGs with most contribution) and decremental (removing COGs with the least contribution) approaches to construct such a set (see Methods for details) only achieved *R*^2^ 0.43 (p-value 2×10^−15^) and *R*^2^ 0.41 (p-value 7×10^−15^) respectively, producing disjoint COG sets, meaning that there is no unique, coherent combination of COGs that would explain the differences in the observed rearrangement to flux ratios (AdditionalData_5; AdditionalData_6).

## Discussion

Evolution of bacterial and archaeal genomes is highly dynamic involving extensive loss and gain of gene, primarily, via HGT, and different types of genome rearrangement. Long range gene order, above the operon scale, is poorly conserved, with little synteny conservation observed even between closely related bacteria and archaea. However, the contributions of different types of processes to this fast decay of synteny in the evolution of prokaryotes have not been quantified. Intuitively, it might seem that intra-genome translocations should be the principal driver of the genome synteny decay. However, gene gains and losses in themselves also break synteny, and moreover, it has been shown that in archaea a gene’s propensity for involvement in genome rearrangement is proportional to its propensity for being gained or lost (25). Therefore, we were interested to quantify the relative contribution of genome rearrangement and gene flux (combined gains and losses) to the genome synteny disruption in prokaryote genome evolution. To this end, we took advantage of the jump model that rigorously derives the SBL distribution from the number of translocations (26) and compared that distribution to the those observed for groups of closely related bacterial and archaeal genomes (ATGCs), for which we also measured the gene content distances reflecting the gene flux. This analysis allowed us to estimate the genome rearrangement to gene flux ratio *q*^∗^ which was found to be robust for the majority of the ATGCs.

Although the overall distribution of the *q*^∗^ values was broad, for the majority of the ATGCs, it fell with the interval of 5-13 gain-loss pairs per translocation. Thus, perhaps, unexpectedly, gene gains and losses seem to account for 85-95% of the genome synteny decay in most bacteria and archaea at the typical ATGC evolutionary distances (<< 1 fixed substitutions per nucleotide). In other words, it appears that, at least at short evolutionary distances, genome dynamics can be largely reduced to gene flux. This finding seems particularly notable because gene gain and loss have been found to occur primarily in hotspots, regions occupying a relatively small fraction of the genomes (6, 31). Gene insertion and elimination within such hotspots has limited effect on synteny. However, apparently, even the occasional gene insertions and losses across the rest of the genomes suffice to cause extensive synteny disruption.

Genes gains and losses seem to be the major cause of synteny loss in prokaryote evolution, but they are clearly insufficient to explain the patterns observed by genome comparison. Gains and losses of individual genes or small gene arrays disrupt synteny blocks but not the overall gene order that produces a diagonal pattern on traditional dot-plot representations of genome comparisons (Figure 4_genome comparison dot-plots) (3, 14). However, with increasing evolutionary distance, this pattern is quickly disrupted, and not only by single-gene translocations addressed here, but even more drastically, by segmental translocations and inversions. The contribution of these larger scale rearrangements is reflected in the deviation of the observed SBL distributions from the jump model expectations, with its pronounced excess of relatively long synteny blocks (Figure 2_SBLcomparison). Apparently, both preservation of synteny in segments consisting of functionally coherent genes (25), and non-random location of gene indels and rearrangements (6) substantially contribute to the overall evolution of the genome organization in prokaryotes. A more realistic theoretical framework for analysis of these different processes of prokaryote genome evolution remains to be developed.

**Figure 4.**
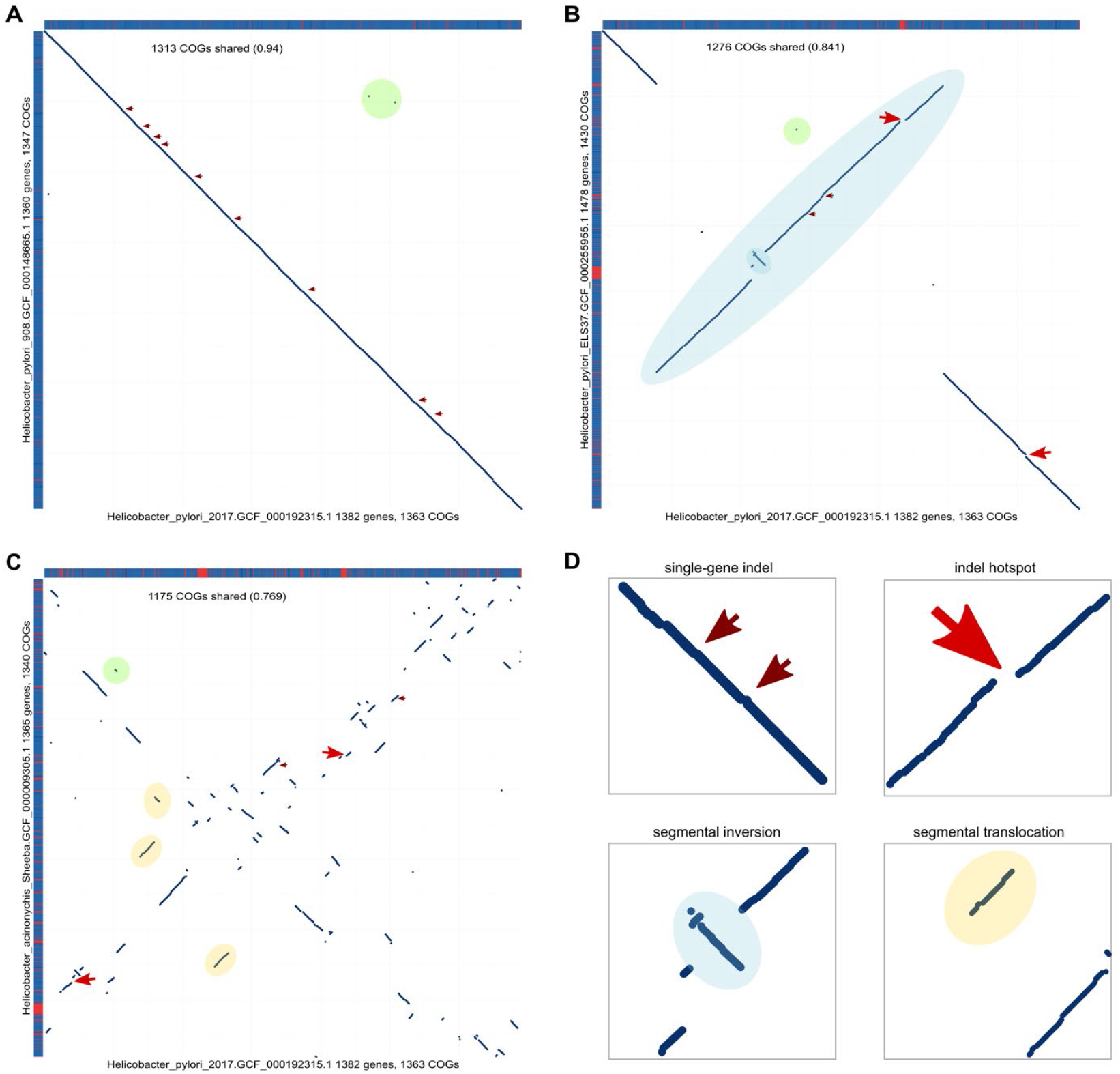
Genome comparison dot-plots for different evolutionary distances. Each dot corresponds to a pair of orthologous genes in the compared genomes. The sides of the panels depict genome composition (blue, shared gene; red, unique gene). Dots indicate locations of orthologous genes. Genomes belong to ATGC050. A. *Helicobacter pylori* 2017 (GCF_000192315.1) vs *H. pylori* 908 (GCF_000148665.1). Evolution of gene order is dominated by single-gene indels (thin red arrows) and single-gene translocations (green circles). B. *H. pylori* 2017 (GCF_000192315.1) vs *H. pylori* ELS37 (GCF_000255955.1). Evolution of gene order is dominated by indel hotspots (thick red arrows) and segmental inversions (blue ellipses). C. *H. pylori* 2017 (GCF_000192315.1) vs *H. acinonychis* Sheeba (GCF_000009305.1). Evolution of gene order is dominated by indel hotspots (thick red arrows), segmental inversions (blue ellipses) and segmental translocations (orange rectangles). D. The inset shows different zoom-in for different types of genome rearrangement events.

## Methods

### Genomic data

A collection of Alignable Tight Genomic Clusters (ATGC) was consists of groups of (nearly) completely sequenced, closely related bacterial and archaeal genomes that were selected for both high sequence similarity and synteny preservation (27). The ATGC dataset includes classification of all protein-coding genes into clade-specific clusters of orthologous genes (ATGC COGs). Distribution of ATGC COGs along the genome partitions (chromosomes and plasmids) provides the intra-ATGC reference for gene order. ATGCs with 3 or more genomes (excluding the largest group, ATGC001, that consists of 432 genomes of *Escherichia, Salmonella* and related *Enterobacteriaceae*) were analyzed; cliques of genomes with identical complements or ATGC COGs were reduced to a single representative.

### Gene content comparison

Gene content distance (GCD) between genomes *A* and *B* within an ATGC is calculated as

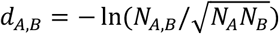

where *N*_*A*_ and *N*_*B*_ are the number of distinct ATGC COGs in the genomes *A* and *B*, respectively, and *N*_*A,B*_ is the number of distinct ATGC COGs shared between these two genomes.

### Gene order comparison

Synteny between two genomes *A* and *B* within an ATGC assessed as follows. First, both genomes were reduced to their common gene complement by eliminating genes in genome *A* that belong to ATGC COGs not represented in genome *B* and vice versa. Then, all synteny blocks (consecutive strings of the same ATGC COGs, regardless of direction, including degenerate blocks consisting of one gene), were identified for this genome pair. Gene order similarity for a pair of genomes was assessed from the SBL distribution. The simplest measure is the mean SBL that ranges from the entire genome length for a pair of identical genomes (all genes form one syntenic block) to one for two completely scrambled genomes (no pair of genes that are adjacent in one genome is adjacent in another, so all syntenic blocks have length one).

### Single-gene neutral rearrangement model

The single-gene neutral rearrangement model (jump model) (26) assumes that gene order evolves by random translocations of a single gene into a random genome location. Evolution under this regime results in predictable decay of synteny with the number of translocations. Specifically, for an asymptotically large linear genome, the fraction of syntenic blocks, retained from the ancestral genome state, is given by:

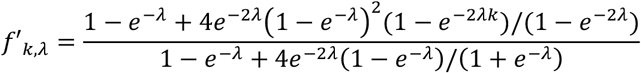

where *f*’_*k,λ*_ is the expected fraction of syntenic blocks of length *k* in the SBL distribution and *λ* is the number of translocations per gene (*k* is a positive integer and *λ* is a finite positive real number).

For any pair of genomes *A* and *B*, real or simulated, an observed SBL distribution 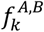 can be compared with the expected SBL distribution *f*’_*k,λ*_ for the given *λ*. The value of *λ* can be optimized to obtain the best fit between 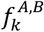 and *f*’_*k,λ*_, yielding *λ*^∗^, the number of genome rearrangements required to produce the given SBL distribution under the single-gene neutral rearrangement model.

### Mapping ATGCs to the Tree of Life

A collection of 47,545 completely sequenced prokaryotic genomes available from GenBank and RefSeq (32) in November 2023 was annotated using COG profiles in the NCBI CDD database(33). Sequences belonging to the set of 54 nearly universal COGs for Bacteria and 55 nearly universal COGs for Archaea were collected and aligned within each COG using FAMSA (34). If more than one paralog per genome was detected, the one most similar to the alignment consensus was used as the index ortholog (35). Alignments were concatenated for Bacteria and Archaea separately and the corresponding trees were reconstructed using FastTree with gamma-distributed site rates and WAG evolutionary model (36).

Genomes from the 139 ATGCs with well-defined *q*^∗^ values were mapped to the leaves of these two trees via NCBI assembly IDs; one random representative per ATGC was selected; the trees were trimmed to these representatives and artificially joined by a Bacteria-Archaea internal branch.

### Analysis of potential genome determinants of the rearrangement-to-flux ratio

First, to determine whether rearrangement-to-flux ratios is correlated with phylogeny, we simulated expected ratio values on the given phylogenetic tree using Brownian motion assumption using *evolvability* package in R (37).

Using the mapping of ATGC genomes to the recent COG-annotated Genomes (38)and the subset of the global phylogenetic tree, representing the relationships between the 139 ATGCs, we analyzed the relationships between the ATGC-specific rearrangement-to-flux ratios and COG-level genome markers.

First, for each COG, its representation in a given ATGC was determined by counting the fraction of genomes within this ATGC that encompassed a representative of that COG, producing a vector of 139 values (mostly, 0 or 1, that is, a COG is represented in either none or in all genomes in an ATGC). Spearman rank correlation between this COG presence vector and the vector of 139 rearrangement-to-flux ratios was calculated and evaluated for robustness via 10,000 random permutations. None of the correlations were both strong (|*r*| ≥ 0.4) and robust (p-value ≤ 0.001).

To account for the phylogenetic information in the data, we utilized phylogenetic generalized least square method with Pagel’s λ as implemented in *caper* package in R {Thomas, 2008 #1911}. Models were built for individual COGs and for COG combinations. The obtained p-values for the individual COG models were corrected with Bonferroni correction for multiple testing. Association between a COG and *q*^∗^ values was considered to be significant if it had a p-value after correction ≤ 0.01.

For COG combinations, incremental (adding COGs with most contribution) and decremental (removing COGs with the least contribution) approaches were used. The incremental approach started from single COG with the lowest Akaike Information Criterion (AIC) value and COGs were incrementally added as long as the model improved, i.e.

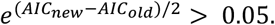

The decremental approach started from the full set of analyzed COGs and gradually removed COGs one by one based on the same criterion. R version 4.2.2 was used for the statistical analysis.

## Data Availability

All the data generated for this work is available in Additional Datasets 1-6 and at https://doi.org/10.5281/zenodo.14806301

## Acknowledgments

S.K.G., Y.I.W, and E.V.K. are funded by the Intramural Research Program of the NIH (National Library of Medicine). S.B. and S.S. were supported by the US/Israel Binational Science Foundation Grant #2021139.

## Author contributions

E.V.K. and S.S. initiated the project and designed research; S.B. and Y.I.W. performed research; S.B., S.K.G. and Y.W.W. analyzed the data; Y.I.W. and E.V.K. wrote the paper that was edited and approved by all authors.

## References

1. Snel B, Bork P, & Huynen MA (2002) Genomes in flux: the evolution of archaeal and proteobacterial gene content. Genome Res 12(1):17–25

2. Kunin V & Ouzounis CA (2003) The balance of driving forces during genome evolution in prokaryotes. Genome Res 13(7):1589–1594

3. Koonin EV & Wolf YI (2008) Genomics of Bacteria and Archaea: the emerging dynamic view of the prokaryotic world. Nucleic Acids Res

4. Koonin EV (2015) The Turbulent Network Dynamics of Microbial Evolution and the Statistical Tree of Life. J Mol Evol 80(5-6):244–250

5. Rodriguez-Valera F, Martin-Cuadrado AB, & Lopez-Perez M (2016) Flexible genomic islands as drivers of genome evolution. Curr Opin Microbiol 31:154–160

6. Wolf YI, Schurov IV, Makarova KS, Katsnelson MI, & Koonin EV (2024) Long range segmentation of prokaryotic genomes by gene age and functionality. Nucleic Acids Res 52(18):11045–11059

7. Trost K, Knopp MR, Wimmer JLE, Tria FDK, & Martin WF (2024) A universal and constant rate of gene content change traces pangenome flux to LUCA. FEMS Microbiol Lett 371

8. Lerat E, Daubin V, Ochman H, & Moran NA (2005) Evolutionary origins of genomic repertoires in bacteria. PLoS Biol 3(5):e130

9. Treangen TJ & Rocha EP (2011) Horizontal transfer, not duplication, drives the expansion of protein families in prokaryotes. PLoS Genet 7(1):e1001284

10. Puigbo P, Lobkovsky AE, Kristensen DM, Wolf YI, & Koonin EV (2014) Genomes in turmoil: quantification of genome dynamics in prokaryote supergenomes. BMC Biol 12:66

11. Mushegian AR & Koonin EV (1996) Gene order is not conserved in bacterial evolution. Trends Genet 12(8):289–290

12. Dandekar T, Snel B, Huynen M, & Bork P (1998) Conservation of gene order: a fingerprint of proteins that physically interact. Trends Biochem Sci 23(9):324–328

13. Wolf YI, Rogozin IB, Kondrashov AS, & Koonin EV (2001) Genome alignment, evolution of prokaryotic genome organization, and prediction of gene function using genomic context. Genome Res 11(3):356–372

14. Novichkov PS, Wolf YI, Dubchak I, & Koonin EV (2009) Trends in prokaryotic evolution revealed by comparison of closely related bacterial and archaeal genomes. J Bacteriol 191(1):65–73

15. Jacob F & Monod J (1961) Genetic regulatory mechanisms in the synthesis of proteins. J Mol Biol 3:318–356

16. Lawrence JG (2003) Gene organization: selection, selfishness, and serendipity. Annu Rev Microbiol 57:419–440

17. Mejia-Almonte C, et al. (2020) Redefining fundamental concepts of transcription initiation in bacteria. Nat Rev Genet 21(11):699–714

18. Lawrence JG (1997) Selfish operons and speciation by gene transfer. Trends Microbiol 5(9):355–359

19. Lawrence J (1999) Selfish operons: the evolutionary impact of gene clustering in prokaryotes and eukaryotes. Curr Opin Genet Dev 9(6):642–648

20. Price MN, Huang KH, Arkin AP, & Alm EJ (2005) Operon formation is driven by co-regulation and not by horizontal gene transfer. Genome Res 15(6):809–819

21. Ballouz S, Francis AR, Lan R, & Tanaka MM (2010) Conditions for the evolution of gene clusters in bacterial genomes. PLoS Comput Biol 6(2):e1000672

22. Eisen JA, Heidelberg JF, White O, & Salzberg SL (2000) Evidence for symmetric chromosomal inversions around the replication origin in bacteria. Genome Biol 1(6):RESEARCH0011

23. Tillier ER & Collins RA (2000) Genome rearrangement by replication-directed translocation. Nat Genet 26(2):195–197

24. Hughes D (2000) Evaluating genome dynamics: the constraints on rearrangements within bacterial genomes. Genome Biol 1(6):REVIEWS0006

25. Wolf YI, Makarova KS, Lobkovsky AE, & Koonin EV (2016) Two fundamentally different classes of microbial genes. Nat Microbiol 2:16208

26. Sevillya G, et al. (2020) Horizontal Gene Transfer Phylogenetics: A Random Walk Approach. Mol Biol Evol 37(5):1470–1479

27. Kristensen DM, Wolf YI, & Koonin EV (2017) ATGC database and ATGC-COGs: an updated resource for micro- and macro-evolutionary studies of prokaryotic genomes and protein family annotation. Nucleic Acids Res 45(D1):D210–D218

28. Galperin MY, Kristensen DM, Makarova KS, Wolf YI, & Koonin EV (2019) Microbial genome analysis: the COG approach. Brief Bioinform 20(4):1063–1070

29. Givens CR & Shortt RM (1984) A class of Wasserstein metrics for probability distributions. Michigan Mathematical Journal 31:231–240

30. Sharda M, Badrinarayanan A, & Seshasayee ASN (2020) Evolutionary and Comparative Analysis of Bacterial Nonhomologous End Joining Repair. Genome Biol Evol 12(12):2450–2466

31. Oliveira PH, Touchon M, Cury J, & Rocha EPC (2017) The chromosomal organization of horizontal gene transfer in bacteria. Nat Commun 8(1):841

32. Sayers EW, et al. (2025) Database resources of the National Center for Biotechnology Information in 2025. Nucleic Acids Res 53(D1):D20–D29

33. Wang J, et al. (2023) The conserved domain database in 2023. Nucleic Acids Res 51(D1):D384–D388

34. Deorowicz S, Debudaj-Grabysz A, & Gudys A (2016) FAMSA: Fast and accurate multiple sequence alignment of huge protein families. Sci Rep 6:33964

35. Wolf YI, Carmel L, & Koonin EV (2006) Unifying measures of gene function and evolution. Proc Biol Sci 273:1507–1515

36. Price MN, Dehal PS, & Arkin AP (2010) FastTree 2--approximately maximum-likelihood trees for large alignments. PLoS One 5(3):e9490

37. Bolstad GH, et al. (2014) Genetic constraints predict evolutionary divergence in Dalechampia blossoms. Philos Trans R Soc Lond B Biol Sci 369(1649):20130255

38. Galperin MY, et al. (2025) COG database update 2024. Nucleic Acids Res 53(D1):D356–D363

