## Supplemental Information APpendix for "Evolution of gene order in prokaryotes is driven primarily by gene gain and loss"

PNAS

### Supporting Information for
#### Evolution of gene order in prokaryotes is driven primarily by gene gain and loss.

Shelly Brezner, Sofya K. Garushyants, Yuri I. Wolf, Eugene V. Koonin and Sagi Snir

Eugene V. Koonin


**This PDF file includes:**

Figures S1-S2

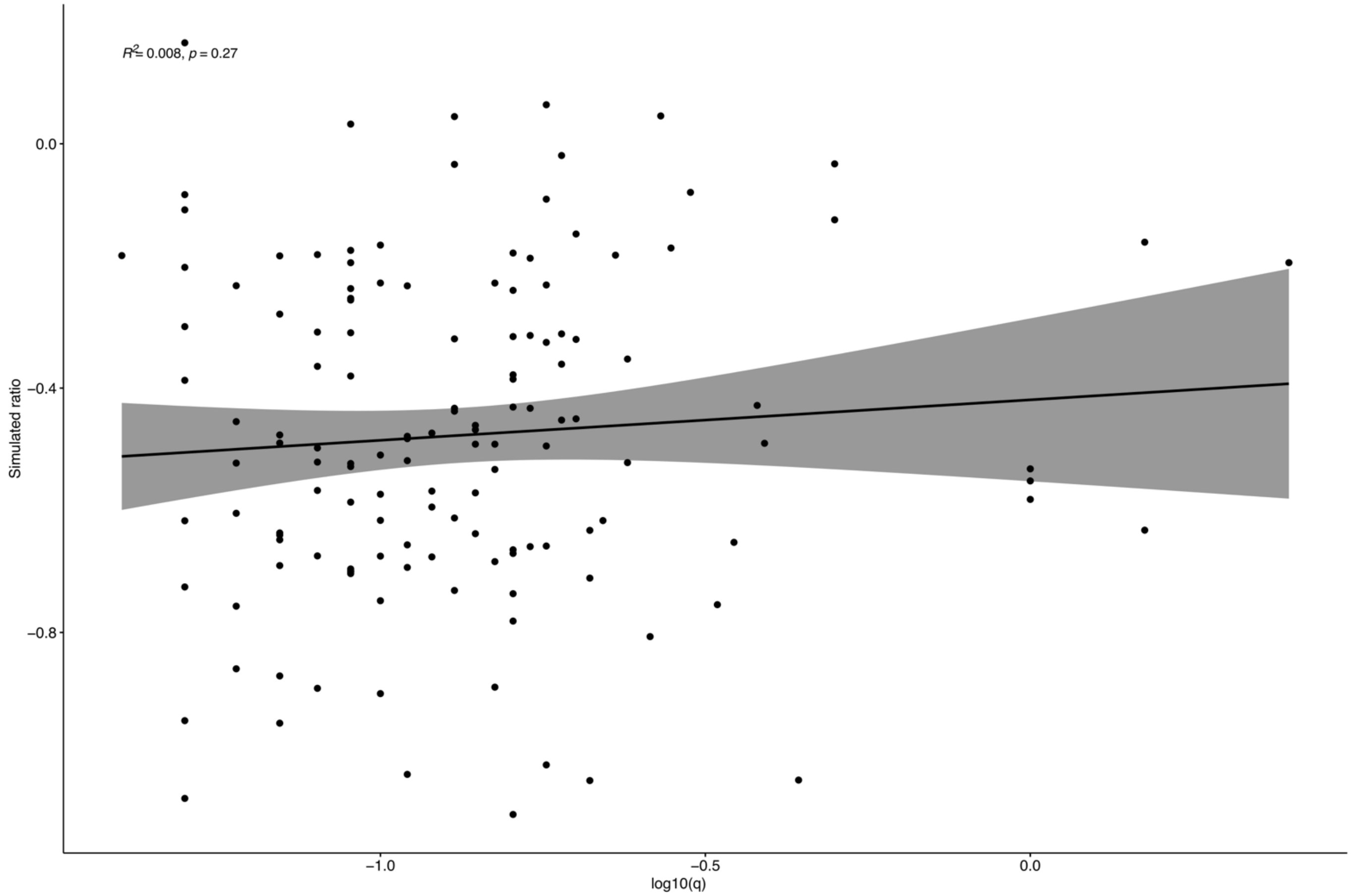


**Fig. S1.** The phylogenetic relatedness between ATGCs is weakly correlated with the rearrangement-to-flux ratio.

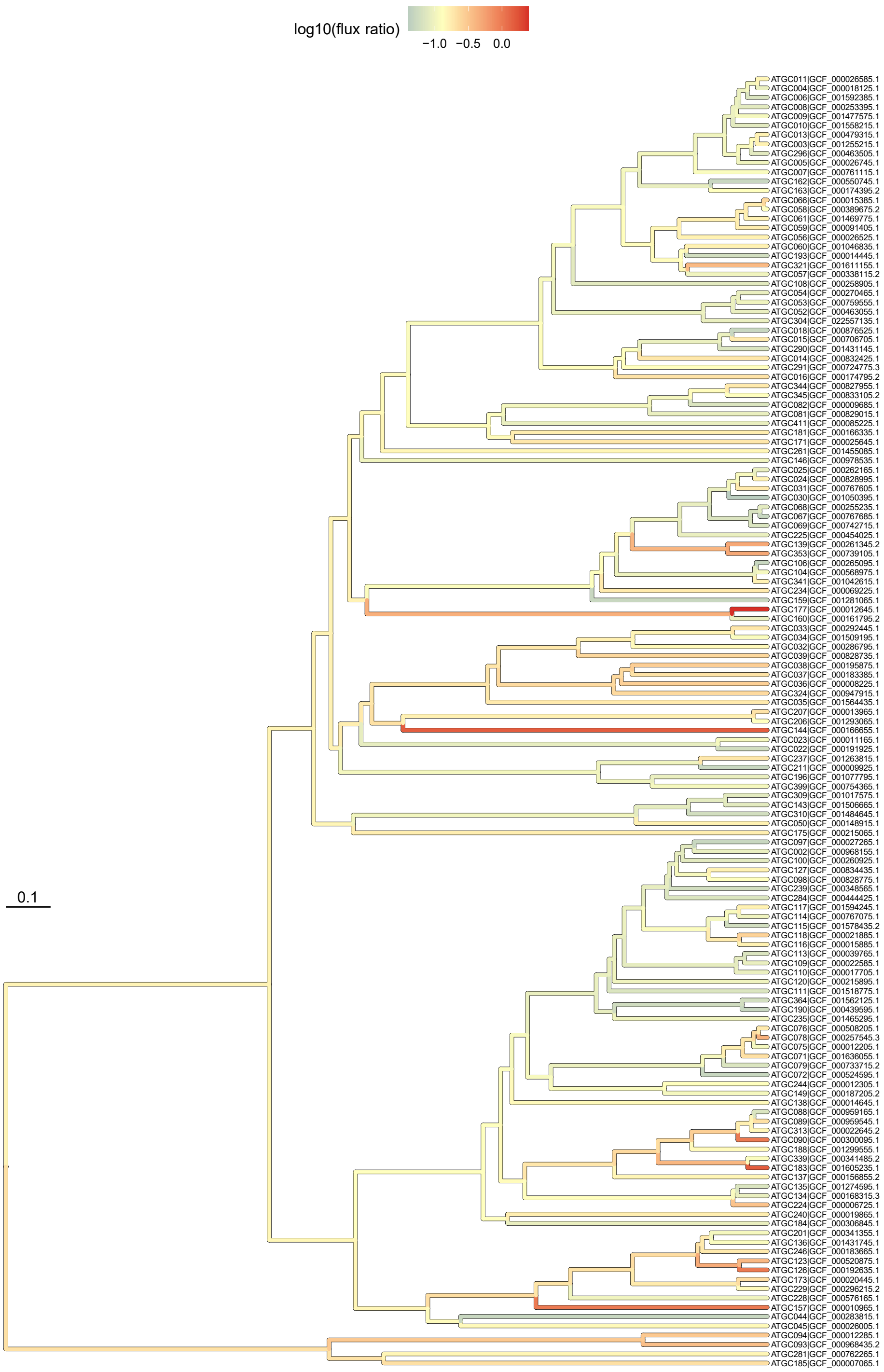


**Fig. S2.** Phylogenetic relationships between the ATGCs with the rearrangement-to-flux ratio ($q^*$) mapped to the tree branches.
